# Intrinsic reinstatement of induced oscillatory context

**DOI:** 10.1101/2021.01.25.428096

**Authors:** Yoann Chantrel, Valeria Trabattoni, Lydia Orton, Amir-Homayoun Javadi

**Author notes:** **Corresponding author:** Amir-Homayoun Javadi, Address: School of Psychology, Keynes College, University of Kent, CT2 7NP, Canterbury, Kent, UK. these authors contributed equally (Yoann Chantrel, Valeria Trabattoni, Lydia Orton).

## Abstract

Declarative memory retrieval is thought to rely on the reinstatement, at retrieval, of contextual cues present during encoding, as evidenced in the context and state-dependent literature. Specifically, previous work has shown that reinstating the oscillatory activity present during encoding, at retrieval, is particularly supportive of memory recall. Our study builds on previous findings suggesting that the oscillatory activity present at encoding may be automatically reinstated during retrieval. To explore the roles of consolidation, prefrontal involvement, and frequency specific activity in this process of oscillatory reinstatement, 115 healthy young adults were randomly assigned to one of five conditions. In each condition, transcranial alternating current stimulation (tACS) was administered at either one of two frequencies (60Hz or 6Hz), over the left dorsolateral prefrontal cortex (left-DLPFC, Experimental) or the right primary motor cortex (right-PMC, Control), during learning of written words. This was followed by a retention interval of either 90 minutes or 1 week and a testing phase, during which EEG activity was recorded. Our results showed significant (and frequency specific) oscillatory reinstatement, after stimulation of the left-DLPFC, in the 1-week retention condition. Oscillatory reinstatement effects were non-significant after stimulation of the right-PMC, or in the 90 minutes retention interval condition. Our results highlight that the oscillatory activity induced during encoding is consolidated as context alongside the information and is reinstated intrinsically during retrieval. The implications of our findings for models of human memory, future avenues of research and clinical applications are discussed.

## 1 Introduction

Context plays a decisive role in supporting declarative memory retrieval, as evidenced by early studies of context-dependent learning (Baker et al., 2004; Godden & Baddeley, 1975) which demonstrated that matching the features of the external environment at encoding and at retrieval enhanced memory performance. Later studies found that signals coming from within the body like physiological arousal (McCaugh, 2000), mood (Koster et al., 2010) or neurological activity, (particularly in structures like the hippocampus or the prefrontal cortex) could provide cues similarly supportive of declarative memory retrieval (Ofen et al., 2007). These findings gave rise to models of state-dependent learning, which posits that memory retrieval is more efficient when individuals are in the same state in which they were when the memory was formed (Radulovic, Jovasevic & Meyer, 2017; Roediger, Tekin & Uner, 2017).

Previous work can be summarised under the framework of the encoding specificity principle put forward by Tulving and Thomson (1973) which posits that recall is most likely when the information available at encoding is also available at retrieval. One of the predictions of this model is, therefore, that a cue will be most likely to lead to retrieval of a memory when there is an overlap between the information contained within this cue and the information to be retrieved. A specific type of neural activity, oscillations, also called brainwaves, is thought to be particularly important in providing contextual cues supportive of declarative memory. This hypothesis is supported by previous studies showing enhanced oscillatory activity during memory retrieval (Addante et al., 2011; Manning et al., 2011) and studies showing that higher similarity between oscillatory activity at encoding and at retrieval is associated with higher chances of successful retrieval (Staudigl & Hanslmayr, 2013).

In line with previous work (Tulvin & Thomson, 1973), these findings gave rise to models suggesting that recall is dependent on the reinstatement, at retrieval, of the oscillatory activity which occurred during encoding (Watrous & Ekstrom, 2014). Causal evidence for these models was brought forward by Javadi et al. (2017) who applied transcranial alternating current stimulation (tACS - known to entrain frequency-specific oscillatory activity; Helfrich et al., 2014) to test if reinstating oscillatory activity present during encoding, at retrieval, will result in improved recall.

In this study, tACS stimulation (at a frequency of either 60Hz or 90Hz) was applied over the left-dorsolateral prefrontal cortex (left-DLPFC), during the encoding of written words. At retrieval, participants were either stimulated with the same frequency as during encoding, or another frequency, while they completed a word recognition task. Recognition was significantly improved when frequencies matched, as opposed to when they did not, providing causal evidence of the role of oscillatory activity in the neural mechanism underlying the effects of state-dependent memory in declarative memory retrieval. Another study of oscillatory reinstatement, conducted by Wimber and colleagues (2012), is of substantial interest to the current study. In Wimber’s study, oscillatory activity was induced at encoding by asking participants to memorise words on a background flickering at either 6 or 10Hz. At retrieval, oscillatory activity was found to re-emerge, at a frequency specific to the one entrained during the encoding of the word retrieved, as measured via electroencephalography (EEG). This suggests that oscillatory reinstatement, found to be supportive of declarative memory retrieval in past studies, may occur naturally at retrieval. These findings also suggest that patterns of oscillatory activity present at encoding must be stored and bound with specific memories during encoding so that they can be simultaneously reactivated during retrieval.

Our study aimed to extend the findings of Wimber et al. (2012) by exploring several aspects of oscillatory reinstatement left unanswered in their study, and respond to concerns raised by Price & Johnson (2018) regarding the method used to entrain oscillation in Wimber et al. (2012), judged too explicit. First, we chose to use tACS to entrain oscillatory activity during encoding, as previous research showed it to be effective in entraining frequency-specific oscillatory activity (Helfrich et al., 2014) and demonstrated that stimulation frequencies cannot be distinguished from one another by participants (Javadi et al., 2017), reducing likelihood of confounding influence. Second, we chose to target the left dorsolateral prefrontal cortex for stimulation, as this area is thought to play a role in the processing of contextual cues (Blumenfeld et al., 2011) and previous research has shown successful modulation of declarative memory via electrical brain stimulation of this area (Javadi, Cheng & Walsh, 2012; Javadi & Walsh, 2012, Zwissler et al., 2014). By stimulating a control site, the primary motor cortex (PMC, chosen for its relative lack of involvement in declarative memory processing), we also tested model-specific predictions of neocortical involvement in oscillatory reinstatement (Watrous & Ekstrom, 2014). Third, in line with previous work suggesting that consolidation supports the integration of contextual cues in memory (McGaugh, 2000) and linking oscillatory activity with this process (Crowley & Javadi, 2019; Javadi & Chang, 2013), we looked at the role of consolidation in the process of oscillatory reinstatement. Specifically, we expected that a longer retention interval would allow for the oscillatory activity induced at encoding to be bound with word items in memory, and integrated, through consolidation, within neocortical networks (Born & Wilhelm, 2012) where the activity linked to their retrieval, could be measured via EEG. Finally, we chose to use two different frequencies (Gamma – 60Hz and Theta – 6Hz), previously shown in the literature to be particularly supportive of declarative memory processes (Staudigl, & Hanslmayr, 2014), to see if oscillatory reinstatement is specific to either one of these frequency bands. Prior work suggests that gamma waves may be more selectively supportive of lower-level perceptual processes like feature binding at encoding (Fell, Fernandez, Klaver, Elger, & Fries, 2003) and theta waves of higher-level binding of distributed cortical memory representations during retrieval (Guderian & Düzel, 2005). However, both gamma and theta frequency-bands have been shown to contribute, independently via phase synchronisation and conjointly via cross frequency coupling, to promoting long-term potentiation and support both declarative memory encoding and retrieval (Nyhus & Curran, 2010; Parish, Hanslmayr & Bowman, 2017).

To test these hypotheses, we asked participants to remember 90 neutral words, while being either stimulated at 60Hz (gamma) or 6Hz (theta). Then, participants were given a retention interval of either 90 minutes or one week. Participants were then tested on a recognition task, while EEG brain activity was recorded.

We hypothesised that we will observe an increase in the specific oscillatory activity induced during encoding, at retrieval, illustrating oscillatory reinstatement. Additionally, we hypothesised that oscillatory reinstatement will be found only after stimulation of left-DLPFC and not the right-PMC. Finally, we expected higher frequency congruent oscillatory reinstatement in the 1-week retention interval condition, than in the 90 min one, consistent with the suggested involvement of consolidation in the binding of contextual cues with memory items in long-term memory.

## 2 Method

### 2.1 Participants

We tested a sample of 113 participants in five testing groups. Participants were randomly assigned to one of five conditions, depending on the type of stimulation (Gamma, 60Hz and Theta, 6Hz), the retention interval (90 minutes and 1 week) and the region stimulated (Experimental and Control groups). Participants were naive to the purpose of the study and the specific condition in which they were in. See Table 1 for a breakdown of the participant demographics in each of the conditions. Participants were right-handed, fluent in English, healthy, with a normal or corrected-to-normal vision, without history of neurological problems, and not currently taking any medications. Participants all gave written informed consent and were all treated in accordance with the Declaration of Helsinki and the guidelines approved by the ethical committee of the School of Psychology at the University of Kent.

**Table 1.**
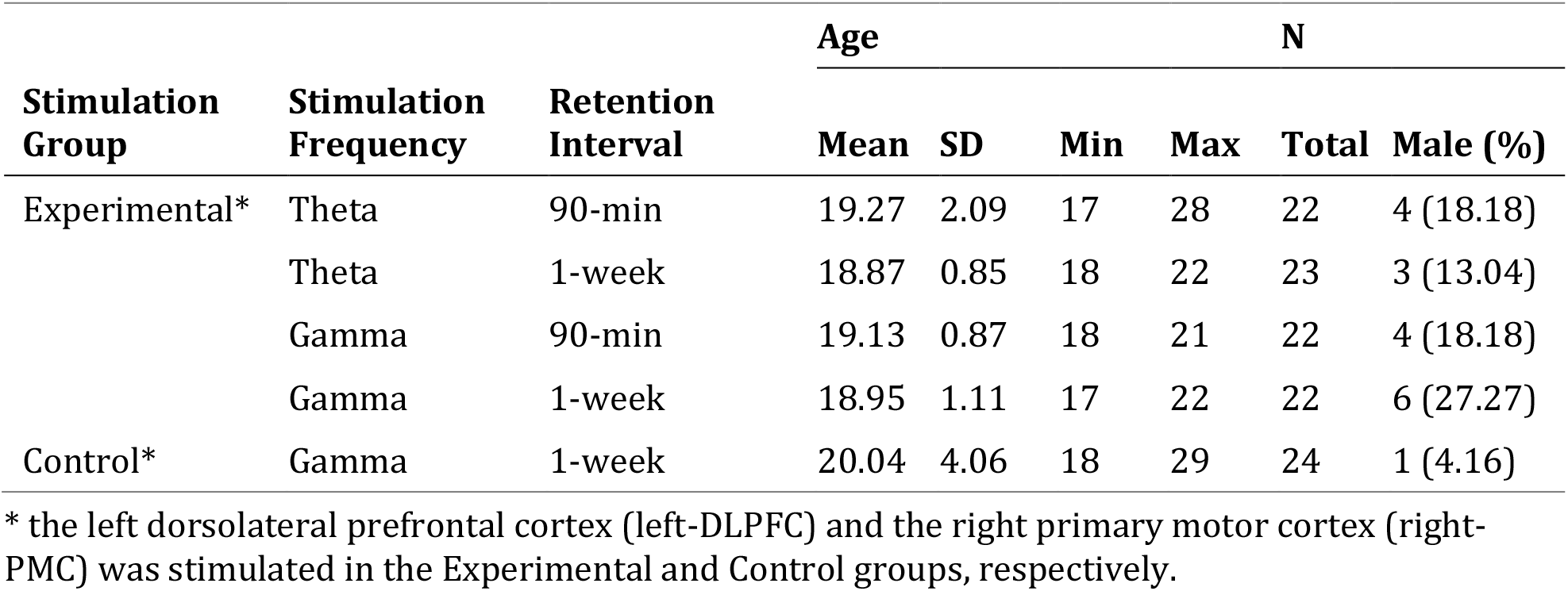
Participants’ demographics by study condition

#### 2.2 Materials

#### Transcranial alternating current stimulation

tACS (neuroConn DC Brain Stimulator Plus, Germany) was administered via two 5×7 cm^2^ saline-soaked surface sponge electrodes. One electrode was placed over the left-DLPFC (Experimental condition; F3 according to the 10-20 international system for EEG electrode placement) or the right-PMC (Control condition; C4), and one electrode on the left wrist. tACS was delivered at encoding with 1.5 mA peak-to-peak amplitude and 1s ramp up and down. Stimulation was delivered at either 60Hz (Gamma frequency) or 6Hz (Theta frequency) depending on the condition. In the Sham and Active conditions, stimulation was delivered for 12 seconds and 10 minutes, respectively. Stimulation began 5 minutes prior to the onset of the stimuli.

#### Stimuli

A bank of 590 words was extracted from The MRC psycholinguistic database (Coltheart, 1981). Words with high valence or similar meanings were excluded. Given that participants were instructed to visualize the words, highly familiar and easily imaginable words were selected. The words were controlled for the number of letters (mean[SD]=4.89[1.24]), number of syllables (1.49[0.50]), printed familiarity (552.54[34.75]), concreteness (580.60[34.06]), and imaginability (581.73[33.20]). For each participant, the words used differed on each day. Words were presented in an Arial font size 18 on a 21-inch monitor, with a 60Hz refresh rate. The stimulus presentation and response recording were conducted using MATLAB v2018 (MathWorks, USA) and the Psychophysics Toolbox v2 (Brainard, 1997).

### 2.3 Procedure

Participants were asked to visualise and memorise words while being electrically stimulated via tACS. They were also told, before learning begun, that they will have to recall the words in a later part of the study. The procedure consisted of an encoding and retrieval phase, separated by a retention interval of either 90 minutes or 1 week, Figure 1. In the encoding part, participants were first given a short practice of the task to get used to its pace. Then, participants took part in the sham stimulation. They were then asked to wait five minutes for the stimulation to ‘come into effect’. Then, while stimulation was meant to be ongoing, participants were shown 45 words, one at a time, (for 0.6s each), followed by an exclamation mark (presented for 4.4s), designating the memorisation period. Once this was completed, the active stimulation part begun and participants were actively stimulated (at either 60 Hz or 6 Hz) for five minutes while waiting. Then, they completed another encoding block of 45 words while stimulation was ongoing. Once this part had been completed, tACS electrodes were removed. Then, participants watched episodes of the Friends TV series for one hour while refraining from using their phones, eating or drinking except water. Once they had done so, participants in the 1-week retention interval left the lab, and came back a week later for the rest of the experiment, while the others took part in the rest of the experiment immediately after the hour had passed. During the final part, at retrieval, all participants were presented with 180 words, divided into two blocks of 90 words, and were asked to decide whether the word presented was old (presented during the encoding phase) or new. While doing this, participants’ brain activity was recorded via EEG.

**Figure 1.**
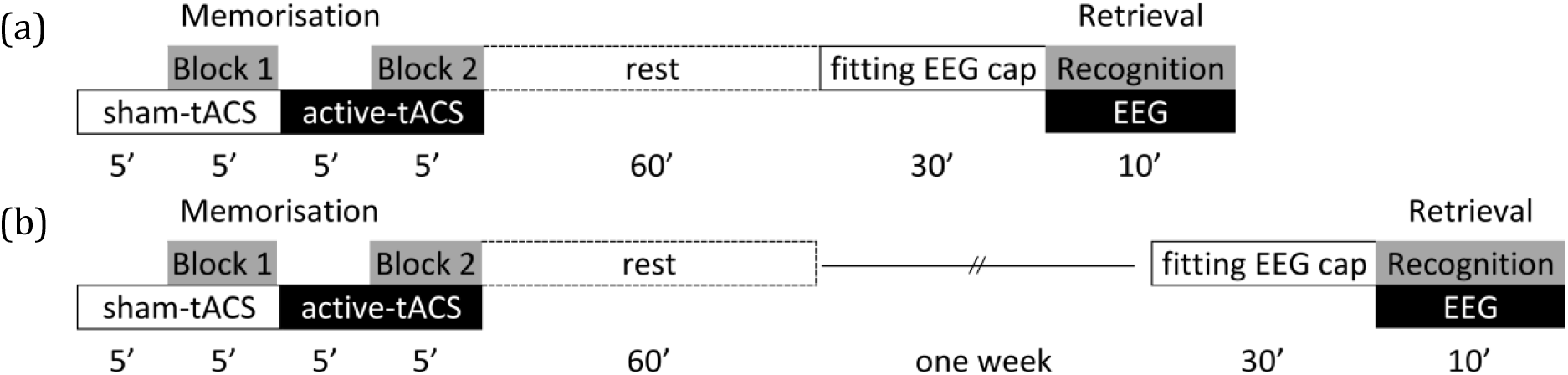
Procedure of the study for (a) 90-min retention interval and (b) 1-week retention interval.

To investigate the site-specificity of the effects of stimulation, a Control group was tested with Gamma-tACS applied to the right primary motor cortex and 1-week retention interval.

#### EEG acquisition and analysis

EEG was set up with a WaveGuard 10/10 layout EEG cap consisting of 32 Ag/AgCl electrode channels (EEGo, Ant Neuro). Position CPz served as reference for Ground. All impedances were kept below 10kΩ. Raw EEG was sampled at 512Hz with 12-bit resolution. EEG data was analysed using SPM v12 (statistical parametric mapping, Wellcome Trust, London, UK). Data was filtered for 0.5-124Hz using 5th order Butterworth filter, montaged based on average electrode activity. Then, eye-blinks were removed using activity of the FP2 electrode and the topography-based artifact correction method: spatial confounds were indicated based on Singular Value Decomposition (SVD) mode and sensor data was corrected using Signal-Space Projection (SSP) correction mode. A maximum of two components of spatial confounds was removed from the EEG data. Subsequently, data was epoched based on -200ms to 1000ms from the moment of the stimulus onset and baseline corrected. Then time-frequency analysis was run using Morlet wavelet transform with 7 wavelet cycles and was scaled based on the pre-stimulus activity to achieve power change. Finally, frequency power data was averaged between 4-8 Hz and between 54-66Hz for Theta and Gamma band activity, respectively. This data was used for statistical analysis as power percentage change of the two frequencies bands, congruent and incongruent to the stimulation frequency. The incongruent frequency was looked at to investigate the frequency specificity of the effects.

Statistical data analysis. SPSS v21 (IBM technologies, USA) was used to carry out statistical analysis. Two mixed-factor analyses of variance (ANOVA) and associated post-hoc tests were conducted. To look at the effect of our independent variables on memory performance, a 2×2×2 mixed-factor ANOVA was conducted, with stimulation condition (old words encoded during Active/Sham stimulation) as a within-participants factor, and stimulation frequency (Gamma/Theta) and retention interval (90-min/1-week) as between-participant factors, and percentage recall of the old words as the dependent variable. To find evidence of oscillatory reinstatement during retrieval of correctly identified old words, a 2×2×2 mixed factor ANOVA was conducted, with stimulation frequency (Theta/Gamma), retention interval (90-min/1-week) as between-subjects factors, and stimulation condition (Active/Sham) as within-subject factors, and power percentage change (changes in frequency power before the onset of stimulus to the onset of stimulus for correctly remembered old words) as a dependent variable. A similar 2×2×2 mixed factor ANOVA was run for incongruent frequency activity. Based on our hypotheses, post-hoc paired t-tests were run to investigate significant interactions.

## 3 Results

### 3.1 Memory performance

The 2×2×2 mixed factor ANOVA investigating the effect of stimulation condition, stimulation frequency, and retention interval on percentage recall, revealed a significant main effect of stimulation condition on memory performance (F(1,109)=19.34, *p*<.001) with higher recall found for old words encoded during sham stimulation (Mean[SD]=66.58[19.42]) than words encoded during active stimulation (59.75[19.76]). A main effect of retention interval was also found (F(1,109)=9.22, *p*=.010), with lower recall found in the 1 week retention interval condition (58.07[16.76]) than in the 90-min retention (68.59[21.77]). All the other interactions or main effects were non-significant, see Table 2 for details. Table 3 summarises the mean and standard deviation for percentage recall in all levels of our independent variables.

**Table 2.** Summary of the results of the mixed-factor ANOVA on percentage recall

| Effect | F | p | $\eta_p^2$ |
| --- | --- | --- | --- |
| Main Effect - Stim. Condition | F(1,109) = 19.344 | <.001 | .151 |
| Main Effect - Stim. Frequency | F(1,109) = 2.648 | .107 | .024 |
| Main Effect - Retention Interval | F(1,109) = 9.224 | .003 | .078 |
| Stim. Frequency × Retention Interval | F(1,109) = 1.645 | .202 | .015 |
| Stim. Condition × Stim. Frequency | F(1,109) = .300 | .585 | .003 |
| Stim. Condition × Retention Interval | F(1,109) = 1.280 | .260 | .012 |
| Stim. Condition × Stim. Frequency × Retention Interval | F(1,109) = 1.971 | .163 | .018 |

Notes: Stim. Condition (old word presented during active/sham stimulation), Stim. Frequency (Gamma/Theta), Retention Interval (90-min/1-week).

**Table 3.** Descriptive statistics of stimulation condition, frequency and retention interval on percentage hit of the old words and percentage correct rejection of the new words (mean[SD]).

| Stimulation Frequency | Retention Interval | New | Old Sham-tACS | Old Active-tACS | t | p | d |
| --- | --- | --- | --- | --- | --- | --- | --- |
| Experimental |  |  |  |  |  |  |  |
| Theta | 90-min | 74.73[16.56] | 67.64[24.54] | 59.27[24.68] | $t(21)=2.467$ | 0.022 | 0.526 |
| | 1-week | 60.83[14.71] | 58.61[17.68] | 55.91[15.54] | $t(22)=1.341$ | 0.194 | 0.280 |
| Gamma | 90-min | 70.73[15.82] | 75.73[15.64] | 71.73[19.33] | $t(21)=1.755$ | 0.094 | 0.374 |
| | 1-week | 55.09[15.09] | 64.18[13.69] | 57.55[12.40] | $t(21)=3.674$ | 0.001 | 0.783 |
| Control |  |  |  |  |  |  |  |
| Gamma | 1-week | 58.21[17.62] | 57.67[19.50] | 54.92[20.32] | $t(23)=1.285$ | 0.211 | 0.262 |

\* d represents Cohen's d effect size.

#### 3.2 Oscillatory reinstatement

A 2×2×2 mixed factor ANOVA was conducted to look at the influence of stimulation condition (Sham-/Active-tACS), stimulation frequency (Theta/Gamma) and retention interval (90 min/1-week) on percentage change in frequency specific power (see Table 4). A significant interaction between stimulation condition and retention interval was found (F(1,85)=12.622, *p*=.001, *η*_*p*_^*2*^=0.129). Paired t-tests were run to test for specific hypotheses. We found a significant difference between power change during retrieval of old word encoded during Active stimulation (143.89[111.62]) compared to Sham stimulation (108.54[87.56]) in the 1-week retention condition (*t*(44)=3.208, *p*=.002, d=0.967). Similar post-hoc comparison for the 90 min retention condition showed a non-significant difference (*t*(43)=1.681, *p*=0.100, *d*=0.512) between Active stimulation (125.20[116.99]) and Sham stimulation (139.20[127.66]) conditions.

**Table 4.** Summary of the results of the mixed-factor ANOVA on power change

| **Effect** | **F(1,85)** | **p** | **$\eta_p^2$** |
| --- | --- | --- | --- |
| Main Effect - Stim. Condition | 2.335 | 0.130 | 0.027 |
| Main Effect - Stim. Frequency | 319.141 | <0.001* | 0.790 |
| Main Effect - Retention Interval | 0.563 | 0.455 | 0.007 |
| Stim. Condition × Stim. Frequency | 0.025 | 0.875 | <0.001 |
| Stim. Condition × Retention Interval | 12.622 | 0.001* | 0.129 |
| Stim. Frequency × Retention Interval | 2.805 | 0.098 | 0.032 |
| Stim. Condition × Stim. Frequency × Retention Interval | 2.324 | 0.131 | 0.027 |

Notes: Stim. Condition (Old words presented during Active-/Sham-stimulation), Stim. Frequency (Gamma/Theta), Retention Interval (90-min/1-week). \* $p < 0.05$ .

To investigate whether the effects of stimulation is frequency specific, we ran a 2×2×2 mixed factor ANOVA was conducted to look at the influence of stimulation condition (Sham-/Active-tACS), stimulation frequency (Theta/Gamma) and retention interval (90 min/1-week) on percentage change in frequency incongruent to the frequency of stimulation in the training phase (Theta for Gamma stimulation and vice versa, see Table 5). None of the effects became significant except main effect of stimulation frequency (F(1,85)=155.842, *p*<0.001, *η*_*p*_^*2*^=0.647).

**Table 5.** Summary of the results of the mixed-factor ANOVA on power change

| **Effect** | **F(1,85)** | **p** | **$\eta_p^2$** |
| --- | --- | --- | --- |
| Main Effect - Stim. Condition | 1.144 | 0.288 | 0.013 |
| Main Effect - Stim. Frequency | 155.842 | <0.001* | 0.647 |
| Main Effect - Retention Interval | 0.978 | 0.325 | 0.011 |
| Stim. Condition × Stim. Frequency | 0.294 | 0.589 | 0.003 |
| Stim. Condition × Retention Interval | 0.192 | 0.662 | 0.002 |
| Stim. Frequency × Retention Interval | 0.041 | 0.840 | <0.001 |
| Stim. Condition × Stim. Frequency × Retention Interval | 0.527 | 0.470 | 0.006 |

Notes: Stim. Condition (Old words presented during Active-/Sham-stimulation), Stim. Frequency (Gamma/Theta), Retention Interval (90-min/1-week). \* $p < 0.05$ .

To investigate that the effects of stimulation is site-specific, we ran a Control condition in which we stimulation right primary motor cortex using Gamma tACS and 1-week retention interval. Comparison of the frequency activity for the Old words presented during Active-tACS (34.40[23.76]) and Sham-tACS (38.34[50.44]) showed a non-significant difference (*t*(23)=0.329, *p*=0.745, *d*=0.067).

See Table 6 for details of the descriptives for different conditions. Figure 2 illustrates power percentage changes in the different levels of our independent variables in the theta and gamma stimulation frequency separately.

**Table 6.**
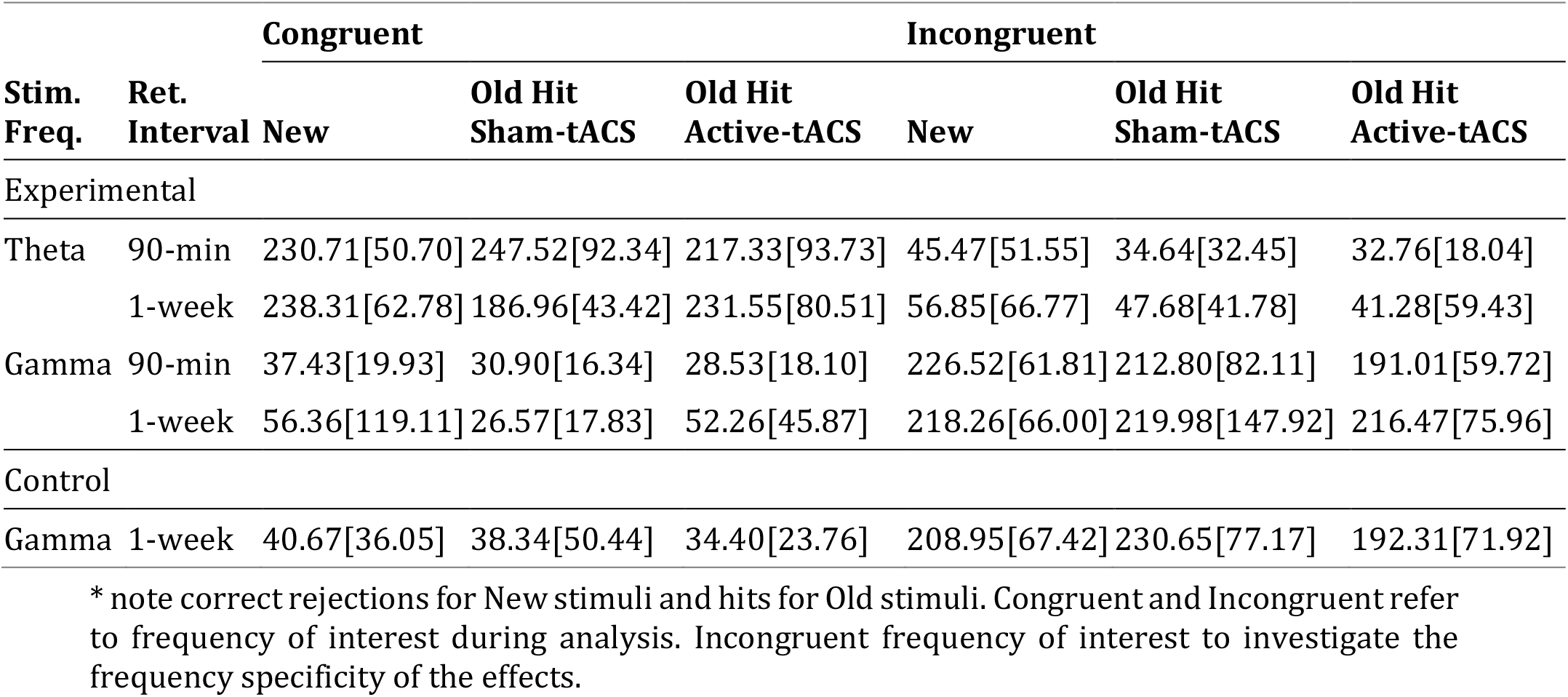
Details of the descriptives for frequency power percentage change during presentation of New words, and Old words memorized during Sham- and Active-tACS.

|  |  | Congruent |  |  | Incongruent |  |  |
| --- | --- | --- | --- | --- | --- | --- | --- |
| Stim. Freq. | Ret. Interval | New | Old Hit Sham-tACS | Old Hit Active-tACS | New | Old Hit Sham-tACS | Old Hit Active-tACS |
| Experimental |  |  |  |  |  |  |  |
| Theta | 90-min | 230.71[50.70] | 247.52[92.34] | 217.33[93.73] | 45.47[51.55] | 34.64[32.45] | 32.76[18.04] |
|  | 1-week | 238.31[62.78] | 186.96[43.42] | 231.55[80.51] | 56.85[66.77] | 47.68[41.78] | 41.28[59.43] |
| Gamma | 90-min | 37.43[19.93] | 30.90[16.34] | 28.53[18.10] | 226.52[61.81] | 212.80[82.11] | 191.01[59.72] |
|  | 1-week | 56.36[119.11] | 26.57[17.83] | 52.26[45.87] | 218.26[66.00] | 219.98[147.92] | 216.47[75.96] |
| Control |  |  |  |  |  |  |  |
| Gamma | 1-week | 40.67[36.05] | 38.34[50.44] | 34.40[23.76] | 208.95[67.42] | 230.65[77.17] | 192.31[71.92] |

\* note correct rejections for New stimuli and hits for Old stimuli. Congruent and Incongruent refer to frequency of interest during analysis. Incongruent frequency of interest to investigate the frequency specificity of the effects.

**Figure 2.**
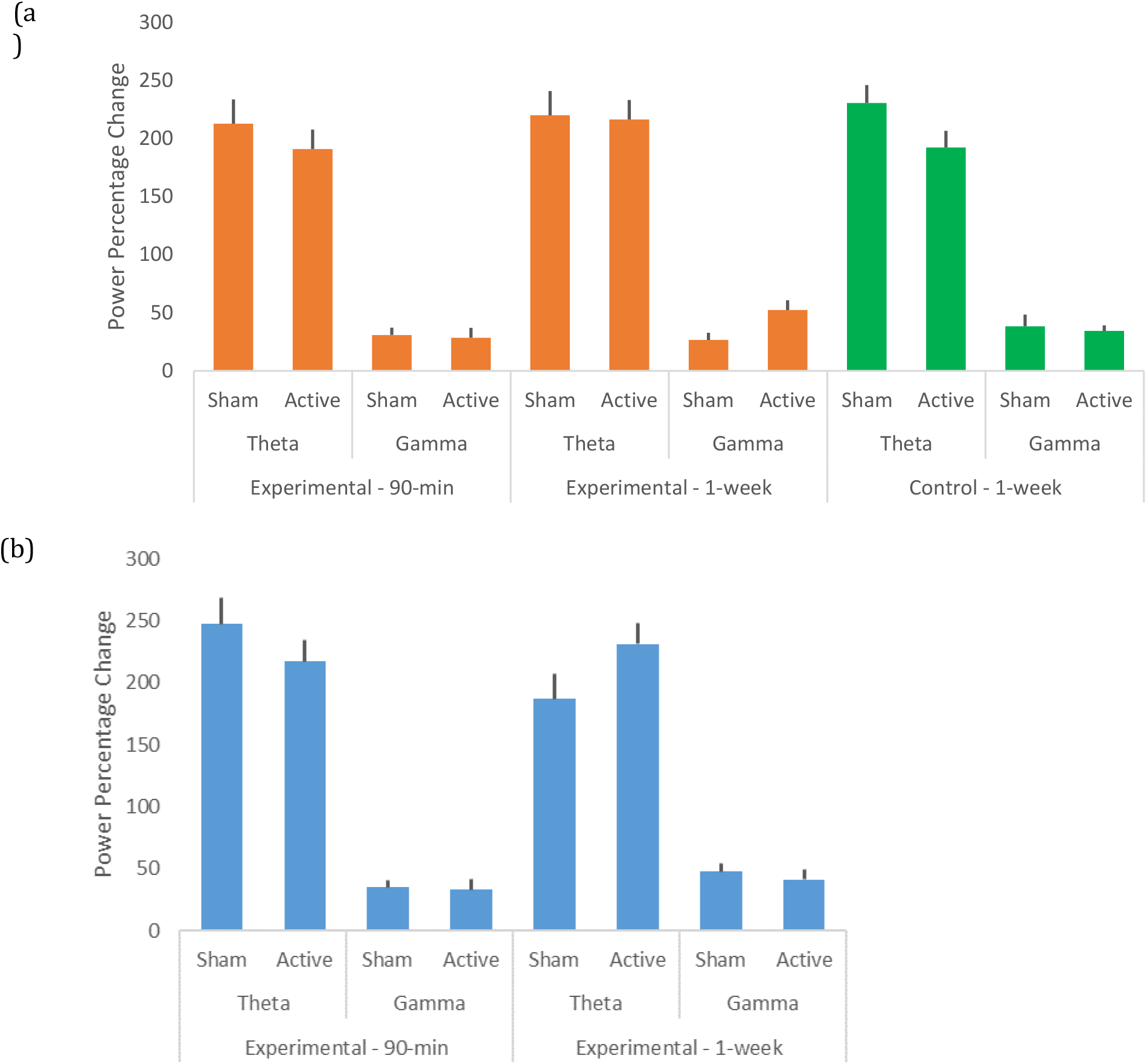
(a) Gamma stimulation frequency, (b) Theta stimulation frequency. Error bars represent one standard error.

## 4 Discussion

In summary, we found lower percentage recall in the 1-week retention condition compared to 90 minutes. We also found higher percentage recall for old words encoded during sham stimulation than for those encoded during active (gamma or theta) stimulation. As hypothesised, we found significantly higher power change (congruent to the oscillatory frequency entrained at encoding via stimulation of the left-DLPFC), during successful retrieval of words encoded during active stimulation, compared to sham stimulation, in the 1-week retention condition. These effects were non-significant in the 90 minutes retention condition. For our control group, who received stimulation over the right-PMC, oscillatory reinstatement effects were non-significant.

Our results come to build on mounting evidence suggesting the importance of oscillatory activity in providing contextual cues supportive of declarative memory retrieval (Staudigl & Hanslmayr, 2014). Declarative memory relies on forming a neural representation of a new experience (which is spatially and temporally grounded within the context in which it occurred) and integrating this representation in a structure which allows for its retrieval based on cues related to the experience itself (Preston & Eichenbaum, 2013). Reinstating the context in which a memory was encoded has been shown to be beneficial to recall, as it provides contextual cues supportive of memory retrieval. This effect was first demonstrated by matching features of the environment at encoding and at retrieval (Godden & Baddeley, 1975, 1980) and later by matching one’s state at encoding and retrieval, primarily via the use of drugs (Kelemen & Creeley, 2003; Young, 2014).

In light of these previous findings, recent neurobiological models have suggested that recall is optimum when patterns of neurological activity established at encoding are reinstated at retrieval (Norman & O’Reilly, 2003; Rugg et al., 2008; Teyler & Rudy, 2007). Specifically, these models posited that oscillatory activity may underpin state-dependent memory (Watrous & Ekstrom, 2014) based on evidence showing that memory retrieval success is associated with the strength of the correlation between neural oscillations during encoding and at retrieval (Sederberg al., 2007; Wimber et al., 2012) and causal evidence showing that matching oscillatory activity at encoding and at retrieval results in enhanced recognition memory (Javadi et al., 2017).

Our findings, which showed superior recognition memory when there was a match between the oscillatory context at encoding and at retrieval (no stimulation for both), relative to a mismatch (stimulation at encoding and no stimulation at retrieval), come to offer direct support to previous models suggesting that oscillatory activity provides contextual cues supportive of memory recall. Specifically, we hypothesise that such supportive effect of oscillatory reinstatement on recall is the result of a mechanism of pattern completion, whereas retrieval cues are transmitted to the hippocampus (particularly the CA3 area), where the pattern of activity created at encoding is ‘completed’ or reconstructed, leading to recall (Rolls, 2013, 2016; Teyler & Rudy, 2007).

However, despite previous studies demonstrating similar effects of oscillatory context on recall (Javadi et al., 2017; Sederberg al., 2007; Wimber et al., 2012) it is worth noting that the superior recall observed for old words encoded during sham, compared to active stimulation, observed in our study, may be due to a primacy effect (Hintzman, Block & Summers, 1973) as the sham condition always preceded the active condition. While this is not the focus of the current study, further work could be done to investigate this potential confounder.

Our findings go further and demonstrate that not only is the reinstatement of patterns of oscillatory activity established during encoding at retrieval beneficial to memory recall, but that this reinstatement may occur automatically at retrieval. Specifically, we showed that inducing frequency specific activity (in two distinct band frequencies; gamma –60Hz and theta –6Hz), during encoding of words, resulted in the reinstatement of congruent oscillatory activity during successful retrieval of said words. Such automatic reinstatement is one of the predictions of the SCERT (Spectro-Contextual Encoding and Retrieval Theory of Episodic Memory; Watrous & Ekstrom, 2014) and is thought to be supported in the brain by phases of synchronisation and de-synchronisation of oscillatory activity, particularly in the theta and gamma frequency bands (Belluscio et al., 2012). Specifically, two oscillatory mechanisms, phase synchronisation (PS) and cross-frequency coupling (CFC) are thought to be involved in coordinating cell assemblies in their initial formation and subsequent reactivation (Axmacher et al., 2010; Canolty & Knight, 2010; Parish, Hanslmayr & Bowman, 2017). PS defines the phenomenon during which activity in one neuronal network reaches its peak in sync with activity in another neuronal network. Alternatively, CFC defines to the phenomenon via which amplitude of fast frequencies is modulated by phases of slow oscillatory activity; CFC manifests itself primarily in gamma amplitude modulated by phases of theta oscillatory activity, also known as theta-gamma cross-frequency coupling (Belluscio et al., 2012). At encoding, theta oscillations influence synaptic activity in the hippocampus via long-term potentiation and depression (Bikbaev & Manahan-Vaughan, 2008) and promote the formation of cell assembly via spike-timing-dependent plasticity (STDP), which relies on synchronised spiking between cells, guided by gamma oscillations (Casula et al., 2016).

Evidence of the role of oscillatory activity in coordinating the formation of cell assembly comes from findings of theta-gamma CFC and gamma-PS increase in the hippocampus, associated with successful memory encoding and enhanced item-context binding (Colgin, 2011, 2015). Moreover, similar oscillatory activity promotes the binding of neocortical and hippocampal cell assemblies in the hippocampus, where they are stored (Parish, Hanslmayr & Bowman, 2017). As such, it is suggested that neural representation of a memory incorporates spatially distributed patterns of oscillatory power (Liu et al., 2009) encoded alongside its engram, and specific to each memory, as evidenced by studies showing that these patterns change depending on the specific item held in memory (Salazar et al., 2012). Studies of oscillatory reinstatement suggested that such a phenomenon occurs within an early time window (within 300ms following stimulus onset) and can involve the reactivation of oscillatory activity unconsciously encoded (Wimber et al., 2012). This suggests that oscillatory reinstatement is part of an early, unconscious, resonance process between a cue (the presentation of a word for instance) and a stored engram, in line with the process of ecphory put forward by Tulving (1983). Taking this previous work into account, in line with previous models (Watrous & Ekstrom, 2014) and evidence of increased CFC and PS during retrieval (Anderson et al., 2009; Düzel et al., 2005), we suggest that, in our study, the presentation of previously encoded words activated an early retrieval signal, which in turn triggered the simultaneous reactivation, (by the hippocampus; Teyler & Rudy, 2007) of neural representations of these specific words and the corresponding patterns of oscillatory activity established in the pre-frontal cortex during their formation (Watrous & Ekstrom, 2014). This, in turn, suggest that oscillations are not only a medium via which brain regions interact, but also a part of a memory, stored alongside engrams, and reactivated together with them at retrieval. While our study focused on activity occurring within the lower theta frequency band and higher gamma frequencies, we suspect that results would not be different within the remaining frequency bands, as they all have been found to be involved in supporting declarative memory processes (Hanslmayr & Staudigl, 2014).

In our study, oscillatory reinstatement was only found after stimulation of the left-DLPFC, and not of the right-PMC. This confirms previous studies showing the importance of the DLPFC in supporting oscillatory reinstatement (Crowley & Javadi, 2019; Javadi & Walsh, 2012; Javadi et al., 2017). Specifically, previous work suggested that the DLPFC is supportive of long-term-memory retrieval by processing the relationship between multiple items in working memory, maintaining and selecting patterns of activity to adapt to task-related demands, and monitoring retrieval cues (Blumenfeld et al., 2011; Braver et al., 2001). Given these findings, is it possible that oscillatory reinstatement was observed because stimulation of the DLPFC lead to better synchrony in neocortical regions, promoting a better integration of contextual information (including oscillatory activity at encoding) within prefrontal networks of contextual cues, leading to the successful retrieval of specific words and simultaneously, of the reactivation of oscillatory activity patterns, encoded alongside their engrams. Given our findings, we support models suggesting that the prefrontal cortex is involved in the processing of temporal context (Polyn & Kahana, 2008) and extend these models to suggest that such temporal contextual cues are expressed as patterns of oscillatory activity and suggest that the left-DLPFC may be involved in the processing and retrieval of such cues. However, while the contrast between the results obtained via stimulation of the left-DLPFC (as compared to our control area, the right-PMC) suggests that involvement of the left-DLPFC at encoding may be necessary for oscillatory reinstatement to occur, it does not mean that left-DLPFC stimulation is sufficient to trigger oscillatory reinstatement. Specifically, due to the limitations inherent to tACS, we could not attribute the effects of our intervention to the stimulation to the left-DLPFC only. Indeed, it has been suggested that tACS can affect regions adjacent to the region of interest (Bikson, Rahma & Datta, 2012), and that its effects may vary based on the individuals’ neurological state at the time of stimulation (Bergmann, 2018). This, along with the dense structural and functional connectivity between the DLPFC and the hippocampus (Anderson et al., 2016; Goldman-Rakic et al., 1984), suggest that it is possible that our findings could have been influenced by activity changes in areas other than the DLPFC, particularly the hippocampus. As such, further work is needed to distinguish the relative contribution of these two areas in the mechanisms of oscillatory reinstatement.

In our study, oscillatory reinstatement was only evident in the participants that had 1 week of retention between encoding (and stimulation of the DLPFC), and retrieval, and not 90 minutes. This suggests that oscillatory reinstatement is dependent on a process called systems consolidation. Complementary learning systems models (Eichenbaum, 2000; McClelland, McNaughton & O’Reilly, 1995; O’Reilly et al., 2014) posit that long-term memory is supported by two specialised systems; the hippocampus which focuses on the rapid acquisition of novel information, and the more distributed neocortex, which focuses on gradually integrating incoming information within structured and contextualised cortical representations (Crowley, Bendor & Javadi, 2019).

Systems consolidation is defined as the process via which memories, at first fragile and dependent primarily on the hippocampus for their storage and retrieval (Teyler & Rudy, 2007) are reorganised in more complex and distributed networks in the neocortex (Smith & Squire, 2009; Squire, Genzel, Wixted & Morris, 2015). This distribution of the activity and the integration of previously encoded oscillatory within a network of contextually related neural representations in the neocortex could be the reason why reinstatement was only found after a 1-week retention interval, due to the limitations of the EEG to measuring only cortical activity.

Alternatively, these findings could illustrate how the integration of contextual information (in this case oscillatory activity at encoding) with memory items (and not only the re-distribution of this information to the neocortex) is time-dependent (Yonelinas et al., 2019). We hypothesise that this integration of contextual information could rely on a mechanism of ‘replay’, thought to occur during consolidation (ólafsdóttir, Bush & Berry, 2018). Previous research demonstrated that declarative memory benefits from the repeated activation of previously encoded engrams (Dudai, 2012; Javadi & Cheng, 2013) particularly during slow-wave sleep (Crowley & Javadi, 2019) and that the hippocampus makes use of time cells (Eichenbaum, 2014) to coordinate the reactivation of previously encoded engrams, following a temporal sequence laid down at encoding (Ji et al., 2007). Following the predictions of the SCERT (Watrous & Ekstrom, 2014), which posit that engrams are responsive to the specific frequencies involved in their formation, it is possible that mechanisms of replay in the hippocampus strengthened encoded oscillatory patterns of activation and promoted their integration with previously encoded memory items, enabling oscillatory reinstatement in the 1 week retention group. However, it is worth noting that performance on our recognition task was close to chance after a 1-week retention interval, so there may have been so undue influence of incorrect retrieval in our findings, which could have lead us to include trials that were misattributed false or true recollection.

Notably, our findings do not fit in with the results of an earlier study conducted by Wimber et al. (2012) who found phase-locked oscillatory reinstatement with a retention interval of only 5 minutes between encoding and retrieval. However, considering a recent failed attempt by Price and Johnson (2018) to reproduce Wimber’s findings, it is possible that oscillatory reinstatement found in their study could be attributed to lenient statistical correction procedures. Alternatively, their results may have been caused by the type of salient and explicit stimulus flicker used to generate oscillation. Indeed, while they have tested for discrimination of flicker patterns and demonstrated that participants were at chance at this task, with a short retention interval like the one used in their study, it is possible that the oscillatory activity observed at encoding and retrieval represents another kind of memory process altogether, driven by unconscious yet neurologically discernible effect of patterned stimulus. However, the lack of significant spatial clusters of oscillatory reinstatement in their EEG analysis does not allow to make good predictions about what that other mechanism could be, so further work could be done to clarify these aspects; for instance, one could try to replace Wimber’s flickering stimulus with a more implicit method for entraining oscillations (like tACS) and keeping a short retention interval, to see if results can then be replicated.

Despite the evident implications of our findings for previous models of memory, we believe that stimulation applied to modulate human memory could have direct clinical applications, as was shown by a recent study demonstrating the beneficial effects of tACS to combat working memory decline in older adults (Reinhart & Nguyen, 2019). A particularly interesting avenue of research lies in the study of schemas. Indeed, in keeping with the early theories of Piaget (1929), evidence has been put forward that schemas are represented in the brain by structured networks of contextually related neural representations, located in the prefrontal cortex, that guides the interpretation of incoming information, and the retrieval of contextually related information by the hippocampus (Gilboa & Marlatte, 2017). Importantly, these schematic networks have been shown to be dysfunctional in multiple clinical presentations, leading to the emergence and maintenance of clinical symptoms (Clark & Beck, 2010; Hitchcock et al., 2017). Based on our findings, which suggest that memories are encoded along with specific representations of oscillatory activity, and that the reactivation of such oscillatory activity can be used to enhance recall for specific items encoded previously (Javadi & Cheng, 2013; Javadi et al., 2017), stimulation could be used to promote the encoding and retrieval of more positive or adaptive material, in an attempt to modify maladaptive schemas in clinical populations.

In conclusion, by manipulating the oscillatory activity during learning of written words (using tACS on the left-DLPFC at two different frequency bands; theta – 6Hz and gamma – 60Hz) and measuring neurological activity during retrieval via EEG, we provided evidence in support of previous models suggesting that oscillatory activity can provide contextual cues beneficial to declarative memory (Watrous & Ekstrom, 2014). Additionally, we demonstrated that retrieval of items encoded during active stimulation lead to the re-activation of the oscillatory activity induced at encoding, a mechanism called oscillatory reinstatement. Specifically, we linked this mechanism to the coordination of cell assembly via mechanism of theta-gamma cross frequency coupling and phase synchronisation (Colgin, 2011, 2015). In our study, such oscillatory reinstatement was only found after a 1-week retention interval following encoding (and not after 90 minutes), we hypothesised that these findings could be explained by the spread of information to the neocortex as put forward in systems consolidation models (Squire, 2004). Alternatively, we posited that these findings could illustrate how the integration of contextual cues relies on a time-dependent process of replay occurring during consolidation (ólafsdóttir, Bush & Berry, 2018). Finally, by also stimulating a control site (right-PMC), we demonstrated that oscillatory reinstatement was found specifically after stimulation of the left-DLPFC, lending support to previous studies showing the importance of this region in oscillatory reinstatement and more widely in supporting declarative memory (Crowley & Javadi, 2019; Javadi & Walsh, 2012; Javadi et al., 2017).

